# Statin-boosted cellular uptake of penetratin due to reduced membrane dipole potential

**DOI:** 10.1101/2020.08.04.236984

**Authors:** Gyula Batta, Levente Kárpáti, Gabriela Fulaneto Henrique, Szabolcs Tarapcsák, Tamás Kovács, Florina Zákány, István M. Mándity, Peter Nagy

## Abstract

Since cell penetrating peptides are promising tools for delivery of cargo into cells, factors limiting or facilitating their cellular uptake are intensely studied. Using labeling with pH-insensitive and pH-sensitive dyes we report that escape of penetratin from acidic endo-lysosomal compartments is retarded compared to its cellular uptake. The membrane dipole potential, known to alter transmembrane transport of charged molecules, is shown to be negatively correlated with the concentration of penetratin in the cytoplasmic compartment. Treatment of cells with therapeutically relevant concentrations of atorvastatin, an inhibitor of HMG-CoA reductase and cholesterol synthesis, significantly increased the release of penetratin from acidic endocytic compartments in two different cell types. This effect of atorvastatin correlated with its ability to decrease the membrane dipole potential. These results highlight the importance of the dipole potential in regulating cellular uptake of cell penetrating peptides and suggest a clinically relevant way of boosting this process.

## Introduction

During the past decades, there has been growing interest in cell penetrating peptides (CPPs), which can traverse biological membranes. CPPs are a diverse set of short peptide sequences that usually consist of 30 or fewer amino acids and can be classified as either cationic, amphipathic or hydrophobic (1). CPPs are important because of their ability to cross cell membranes in a nontoxic manner and because of their capacity to support efficient delivery of cell-impermeable therapeutic cargos with molecular weights several times greater than their own (2). Several natural peptides with cell penetration capability have been characterized including substance P analogs, the Tat protein in HIV and the homeodomain of the Antennapedia protein in Drosophila (3–5). The cell translocation sequence was localized to the third helix of the homeodomain leading to the development of a 16-amino acid oligopeptide rich in positively charged amino acids. This peptide, penetratin, belongs to the cationic class of CPPs, and is widely used in research aimed at defining mechanisms of cellular uptake (6).

While CPPs hold great promise in drug delivery, their clinical potential is currently limited by low bioavailability, short half-life and lack of specificity (7–9). This latter shortcoming can be remedied by equipping the CPP with a homing domain or by fusing it to an inhibitory domain made up of negatively charged residues that is removed in the tumor microenvironment having increased proteolytic activity (9, 10). The mechanism of cellular entry of CPPs also limits their efficiency putting it at the forefront of current investigations. One of the two, well-established routes of cellular entry for CPP-cargos is direct plasma membrane translocation, which may involve formation of inverted micelles, transient pores or increased fluidity of the plasma membrane (2, 11). Another well-established route of cellular entry for CPP-cargos is endocytosis (12). Unless the endocytic uptake itself is followed by endosomal escape, the CPP does not gain access to the cytosolic compartment and is digested in lysosomes. Many studies focused on the release of CPPs from endosomes, leading to the insertion of endosomolytic sequences into or covalent coupling of endosomolytic compounds to CPPs (13, 14).

Numerous other approaches have been adopted to increase the cellular uptake of CPPs including backbone cyclization, unnatural amino acids, pegylation and acylation (14–17). The problem is further complicated by the fact that cellular uptake in 3D tumor spheroids is not strongly correlated with the uptake in monolayers (18). Other strategies for improving cellular delivery are based on the realization that a CPP must cross a membrane independent of its uptake mechanism. The direct translocation mechanism involves crossing the plasma membrane, whereas the endocytic mechanism relies on traversing membranes of the endolysosomal compartment. Due to their charged nature electrostatic interactions of CPPs with anionic phospholipids and heparan sulphate proteoglycans have been implicated in direct membrane translocation and endocytosis, respectively (19, 20). Transport of charged substances across the plasma membrane is also influenced by the three different kinds of membrane potentials, the transmembrane, the surface and the dipole potential (21). The magnitude of the dipole potential, generated by the preferential orientation of lipids and interfacial water molecules, is approximately 200-300 mV, larger by a factor of at least 4-5 than the widely known transmembrane potential (22). Since the electric field associated with the dipole potential is confined to the surface of the membrane, its strength is 10^8^-10^9^ V/m, larger by 1-2 orders of magnitude than the field associated with the transmembrane potential. Therefore, the dipole potential exerts significant effects on the conformation of transmembrane proteins (23–26), on the binding of molecules to the membrane (27) and their transmembrane transport (28). One of the most important factors determining the dipole potential in the sterol content of membranes. Cholesterol has been shown to increase the membrane dipole potential directly due to its intrinsic dipole moment, and indirectly by increasing the order of lipids and interfacial water molecules and by changing the dielectric constant of the membrane (26, 29–32). Due to this correlation the dipole potential has been shown to be larger in raft-like membrane domains in cellular plasma membranes (33). Statins, inhibitors of 3-hydroxy-3-methyl-glutaryl-CoA (HMG-CoA) reductase, decrease the cholesterol content of cells in experimental and clinical settings, and they have also been reported to decrease the dipole potential of the plasma membrane (32, 34). Statins are the most commonly used therapeutic agents to treat hypercholesterolemia due to their beneficial effect on cardiovascular morbidity and mortality (35–38). Although adverse effects, e.g. myopathy, liver dysfunction and type 2 diabetes, have been associated with statins, they are usually well tolerated and successfully used even in combination with other drugs such as cholesterol absorption inhibitors or fibrates (38–41). While the primary mechanism of action of all statins is identical, there are significant differences in their efficacy and bioavailability (40). Atorvastatin is superior to other statins in requiring lower milligram equivalent doses to achieve the same effect on LDL-cholesterol levels (42). As opposed to simvastatin and lovastatin, which are pro-drugs of the active hydroxy-acid form, atorvastatin does not require enzymatic activation, a property not to be overlooked in in vitro applications (43).

Corollary to the aforementioned principles membrane potentials are expected to influence the uptake of penetratin due to its charged nature. However, only a limited number of such studies have been reported. Non-physiological abolishment of the transmembrane potential has been shown to inhibit the uptake of positively charged cell-penetrating peptides (44). Although a negative dipole potential favors the incorporation of cell-penetrating peptides into lipid monolayers in molecular dynamics simulations and in experiments (45, 46), such effects have not been described in lipid bilayers or living cells. Here, we not only show that the physiological, positive dipole potential of cellular membranes inhibits the uptake and endo-lysosomal escape of penetratin, but also report that an artificial decrease of the dipole potential and treatment with atorvastatin in concentrations corresponding to clinical treatments stimulate entry of penetratin into the cytosol.

## Results

### Fluorescence labeling reveals cellular uptake and delayed endo-lysosomal escape of penetratin

Penetratin was labeled with two different fluorescent dyes in order to study the mechanism and kinetics of its cellular uptake. The fluorescence of naphthofluorescein (NF) is quenched at acidic pH enabling studying the release of penetratin from the acidic endo-lysosomal compartment (47). In order to complement the information on endo-lysosomal escape we chose labeling with AFDye532, a dye exhibiting pH-independent fluorescence to report on the total cellular content of penetratin. The cellular fluorescence of AFDye532-penetratin is proportional to the total cellular uptake of penetratin, the intensity of NF-penetratin characterizes its concentration in non-acidic compartments (mainly the cytosol), while the ratio of NF-labeled and AFDye532-labeled penetratin intensities reveals the fractional escape of the cell-penetrating peptide from acidic compartments. The molecular weight and purity of labeled penetratin was checked by mass spectrometry and high performance liquid chromatography (Suppl. Fig. 1). We developed an approach based on flow cytometric measurement of cell-associated fluorescence signals to separately measure the kinetics of cellular uptake and endo-lysosomal release of these fluorescently labeled penetratins. Cells were incubated in the continuous presence of an equimolar mixture of NF-penetratin and AFDye532-penetratin (5 μM of each) at 37°C, and the fluorescence intensity of cells was measured in a time-correlated manner. After removing cells with compromised membrane permeability based on DAPI staining from the dataset and compensating for spectral crosstalk the time-dependent change of fluorescence intensity was plotted using moving average smoothing. The signal of AFDye532-penetratin, corresponding to the total cellular content of penetratin, reached saturation at 200-400 seconds in two different cell lines, while a significantly delayed saturation of penetratin concentration in the neutral, cytoplasmic compartment at 800-1000 seconds was observed based on the pH-sensitive fluorescence of NF-penetratin (Fig. 1). The ratio of NF to AFDye532 intensities, characterizing the fraction of penetratin in neutral compartments, initially declined, corresponding to a preferential presence of penetratin in acidic endosomes, followed by a gradual increase reaching saturation at approximately 800-1000 seconds (Fig. 1). Since flow cytometry lacks subcellular resolution, we correlated the fluorescence intensities of the two reporters with their cellular location. Confocal microscopic investigation of the distribution of the fluorescent penetratin derivatives revealed that AFDye532-penetratin exhibits bright fluorescence even in endosomes where the fluorescence of NF is quenched and that the cell-associated fluorescence of NF originates from outside the endo-lysosomal compartment (Fig. 2). This observation confirms that interpretation of the fluorescence intensity of AFDye532-penetratin and NF-penetratin, measured by flow cytometry, as total cellular penetratin uptake and the amount outside endosomes, respectively, is indeed correct. Since AFDye532 and NF form a Förster resonance energy transfer pair, anticorrelation between their fluorescence intensities could also have been caused by energy transfer, i.e. a high local concentration of the acceptor (NF) quenching the fluorescence of the donor (AFDye532). In order to exclude this possibility, the equimolar mixture of fluorescent penetratins was supplemented with 10 μM unlabeled penetratin. If energy transfer is to blame for the anti-correlation between the fluorescence intensities of the two dyes, dilution of their local concentration with unlabeled penetratin is expected to eliminate or reduce this anti-correlation. However, the time dependence of the fluorescence intensities and their ratio were not changed significantly by the presence of unlabeled penetratin allowing us to conclude that the fluorescence intensities of AFDye532 and NF correctly report total cellular uptake of penetratin and its concentration in neutral compartments, respectively (Suppl. Fig. 2). In summary, we established a flow cytometric approach revealing the different kinetics of cellular uptake and endo-lysosomal release of penetratin setting the stage for further analysis of the effect of dipole potential-modifying agents on penetratin uptake.

**Figure 1.**
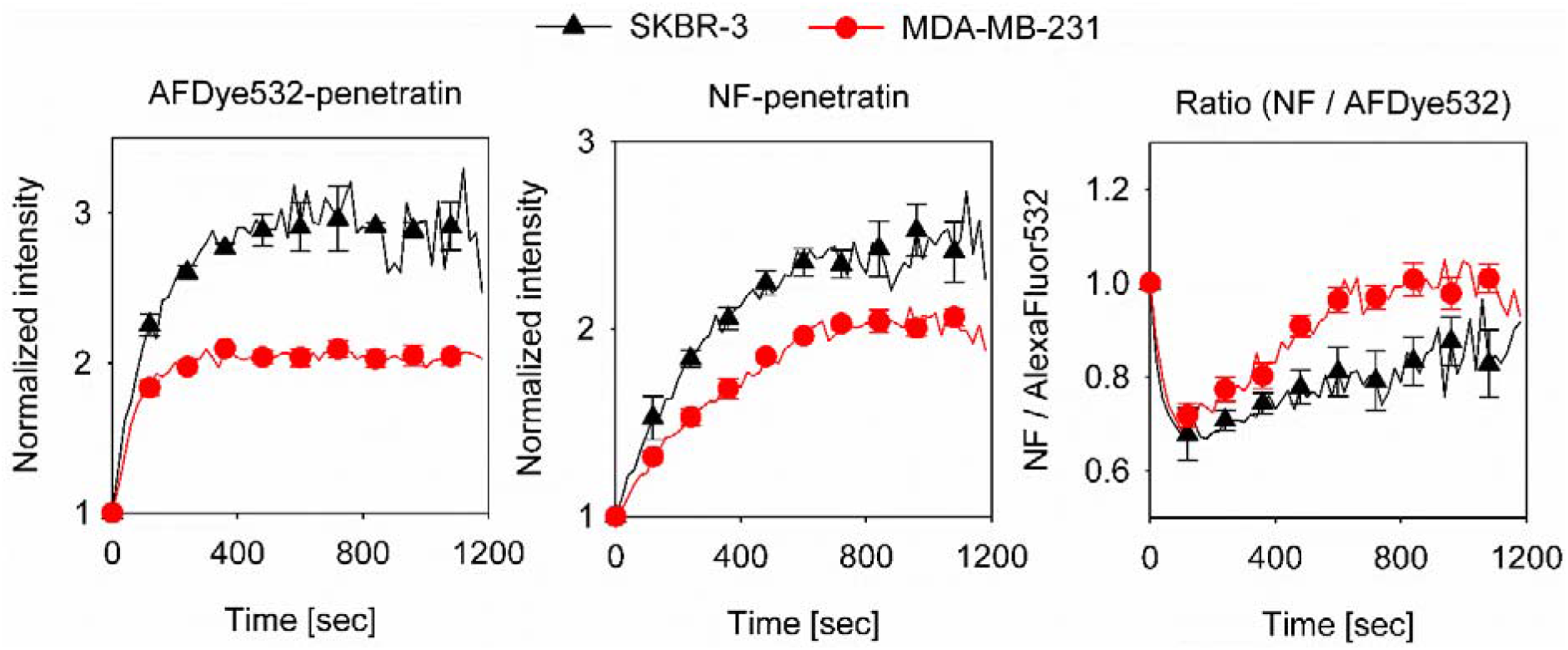
Cellular uptake and endo-lysosomal escape of penetratin. Two different cell types were incubated at 37°C in the continuous presence of fluorescently labeled penetratins (5 μM AFDye532-penetratin and 5 μM NF-penetratin). Time-correlated flow cytometric recording of the fluorescence intensity of cells was performed. The average intensity of AFDye532, characterizing total cellular uptake, and the average intensity of NF-penetratin, characteristic of the amount of penetratin present in pH-neutral compartments, were calculated as a function of time after gating out debris and non-viable cells. The ratio of the two fluorescence intensities is plotted in the panel on the right. The continuous lines show the average fluorescence intensity calculated for 20-sec time periods. The symbols with error bars, representing the standard error of the mean calculated from 10-12 samples from four biological replicates, are only shown for every 7^th^ data point for clarity.

**Figure 2.**
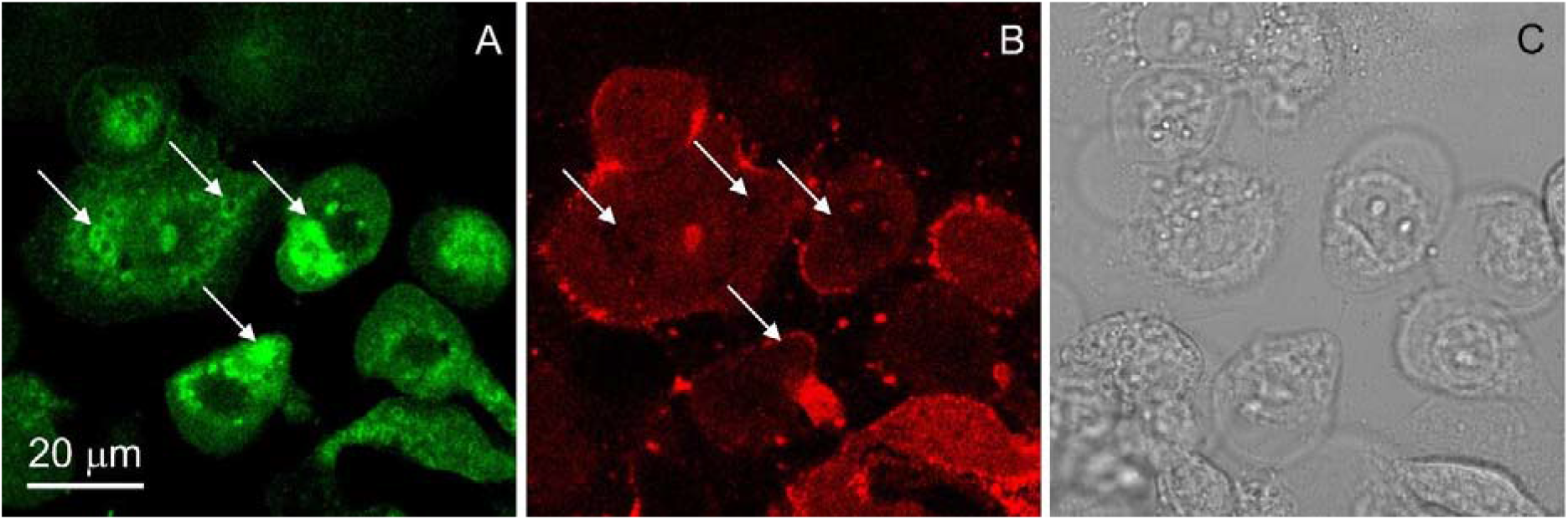
Confocal microscopic images of live cells dual-labeled with AFDye532- and naphthofluorescein-penetratin. Cells were incubated with AFDye532-penetratin (A) and NF-penetratin (B) at 37°C for 20 min and images were captured by confocal microscopy. Panel C displays a differential interference contrast image of the cells. The arrows in the fluorescence images point at areas showing the anti-correlation between the pH-insensitive fluorescence of AFDye532 and the fluorescence of NF quenched at acidic pH.

### Reduction of the membrane dipole potential enhances uptake and endo-lysosomal release of penetratin

Since the strong intramembrane dipole potential is expected to affect the uptake and/or endo-lysosomal release of positively-charged penetratin, we treated cells with 6-ketocholestanol and phloretin, agents known to increase and decrease, respectively, the positive, intramembrane dipole potential (24, 48). Flow cytometric analysis of the fluorescence intensity ratio of the dipole potential sensitive dye, di-8-ANEPPS, confirmed that 6-ketocholestanol significantly increased the dipole potential. On the other hand, phloretin did decrease the dipole potential, but its effect did not reach statistical significance (Fig. 3A). These observations are in accordance with our previous results (24).

**Figure 3.**
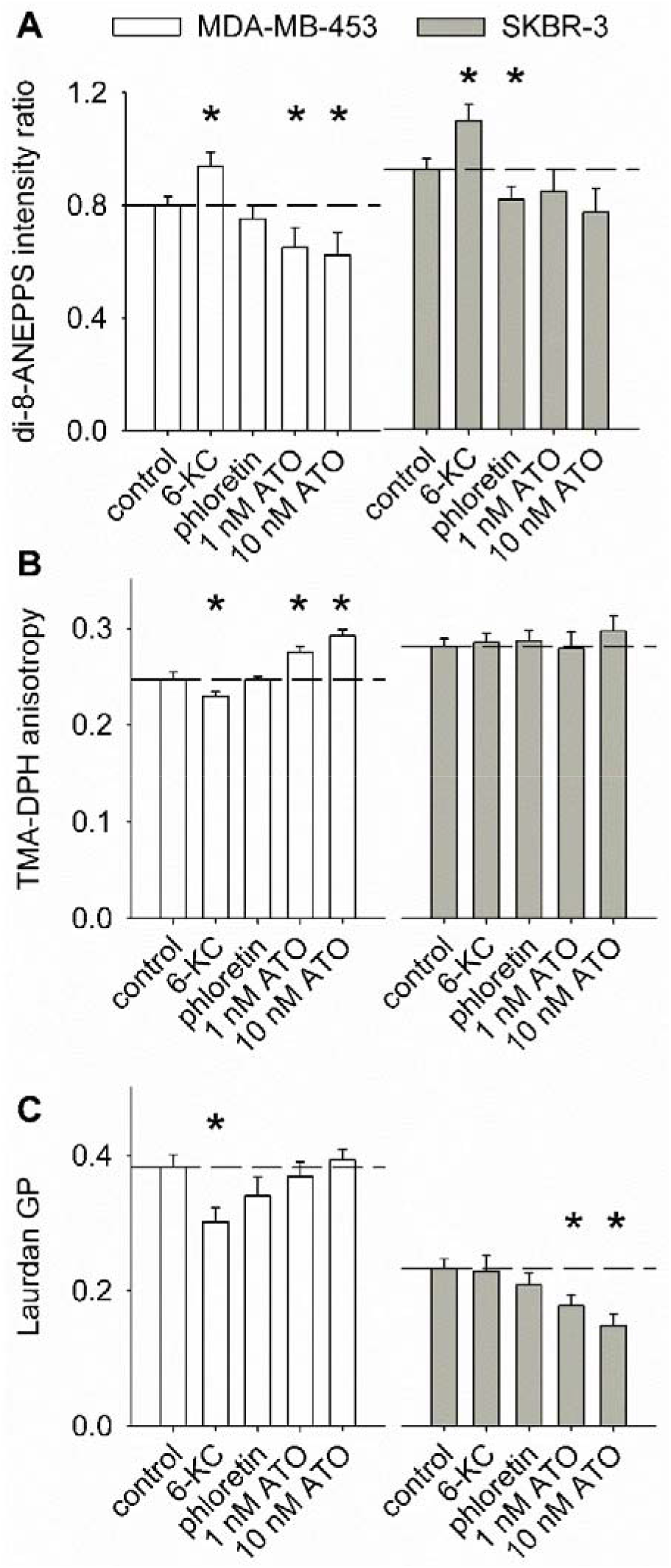
Characterization of the effects of phloretin, 6-ketocholestanol and atorvastatin on the biophysical properties of the membrane. Cells were pretreated with phloretin, 6-ketocholestanol or two different concentrations of atorvastatin (ATO) as described in Materials and methods. The dipole potential was measured with di-8-ANEPPS, whose excitation intensity ratio correlates positively with the dipole potential (A). The fluorescence anisotropy of TMA-DPH is inversely related to membrane fluidity (B), whereas the generalized polarization (GP) of Laurdan is proportional to the hydration or compactness of the membrane (C). The error bars display the standard error of the mean of six independent samples from two biological replicates. Asterisks indicate statistical significance (p<0.05) reported by ANOVA followed by Tukey’s HSD test.

Having established that the treatments modify the dipole potential according to expectations, we set out to characterize the effect of an altered dipole potential on the uptake of penetratin. A diminished dipole potential significantly enhanced both the total cellular uptake of penetratin and its concentration in non-acidic compartments in both SKBR-3 and MDA-MB-231 cells (Fig. 4). Since both of these processes were enhanced at the lower dipole potential achieved by phloretin treatment, the ratio of the NF and AFDye532 intensities, characterizing the fractional release from acidic compartments, remained unchanged. On the other hand, 6-ketocholestanol, which significantly increased the dipole potential (Fig. 3A), had a statistically non-significant effect on both the total cellular uptake and endo-lysosomal release of penetratin (Fig. 4). In conclusion, we have established that the physiological level of the intramembrane, positive dipole potential significantly inhibits the uptake of penetratin.

**Figure 4.**
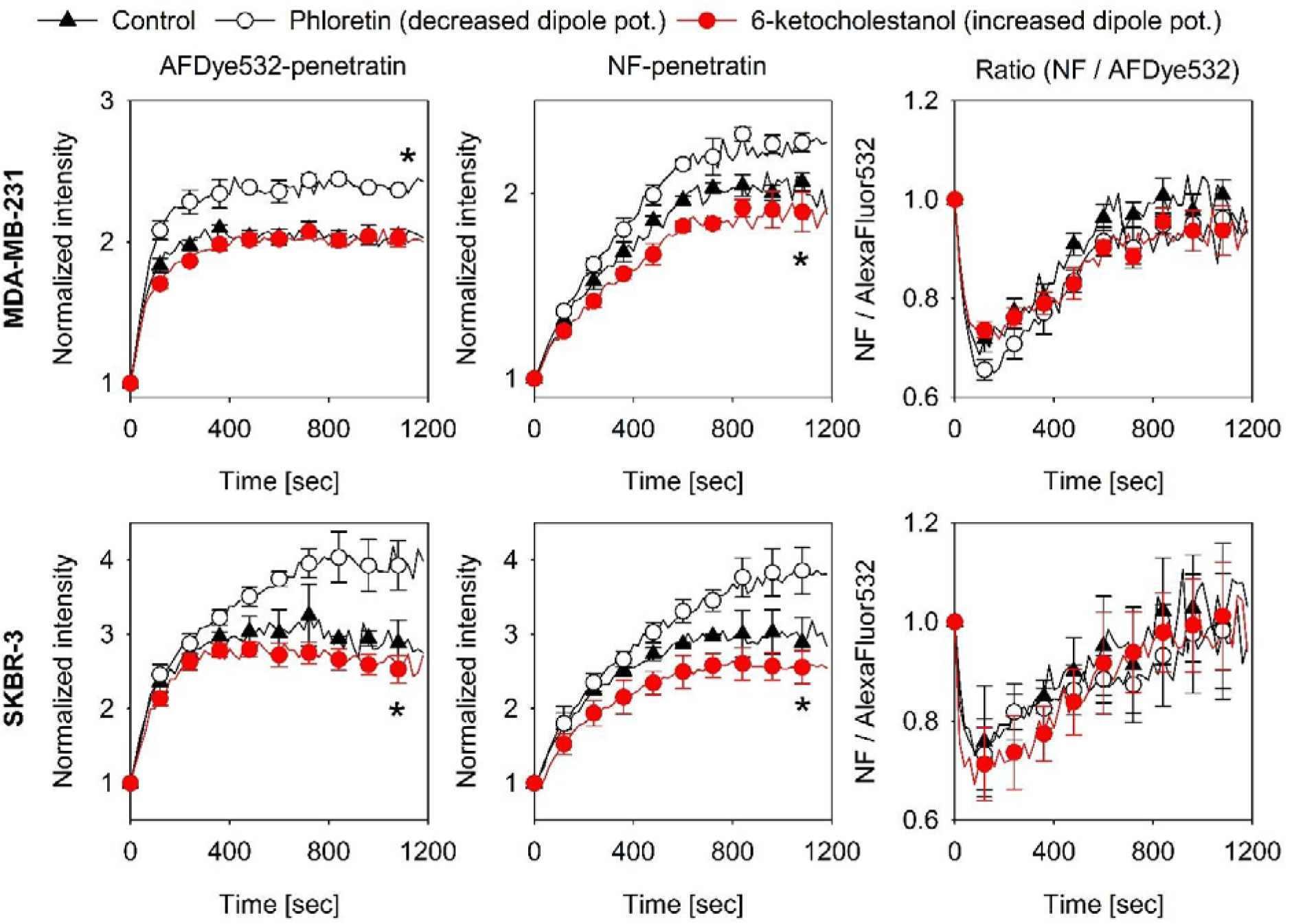
Decreased dipole potential enhances cellular uptake and endo-lysosomal escape of penetratin. Cells (SKBR-3, MDA-MB-231, displayed on the left) were treated with a mixture of AFDye532-penetratin and NF-penetratin at 37°C and their fluorescence intensity was measured by flow cytometry. The membrane dipole potential of cells was decreased and increased by pre-treatment with phloretin and 6-ketocholestanol, respectively. Time-dependent intensities and their ratio were determined after gating out debris and dead cells. The error bars represent the standard error of the mean calculated from 10-12 samples from four biological replicates. Asterisks indicate significant difference of the phloretin-treated sample compared to the control at 20 min (p<0.05).

### Statins increase the endo-lysosomal release of penetratin due to decreased dipole potential

Increasing the uptake of cell penetrating peptides has great potential medical benefit. Since the treatment used for decreasing the dipole potential in the previous section cannot be applied in humans, we sought an alternative approach to enhance the uptake of penetratin by modulating the dipole potential. The dipole potential correlates with membrane cholesterol content (32, 33), and statins are known to decrease the dipole potential (32). We opted for atorvastatin since it is one of the most effective statins and it is the active substance not requiring enzymatic activation (42, 43, 49). Atorvastatin, used at a concentration of 1-10 nM corresponding to the serum concentration in human patients (34), significantly decreased the total cholesterol content of MDA-MB-231 cells by 40-50%. Albeit to a lesser extent, atorvastatin also decreased the cholesterol content of SKBR-3 cells (Suppl. Fig. 3). In perfect agreement with these results atorvastatin decreased the dipole potential in both cell lines, but SKBR-3 displayed lower sensitivity (Fig. 3A). Treatment of MDA-MB-231 cells with the same concentration range of atorvastatin significantly enhanced the endo-lysosomal release of penetratin with only a marginal effect on total cellular uptake (Fig. 5). At the same time, the effect of atorvastatin on SKBR-3 cells was smaller and it did not reach statistical significance. Since the extent of decrease in the cholesterol content of this cell line turned out to be smaller compared to MDA-MB-231, we also tested the effect of higher atorvastatin concentrations. 100 nM and 10 μM of atorvastatin decreased the total cellular uptake of penetratin. Although cells with increased membrane permeability were discarded from the analysis, we attribute this finding to compromised cell viability. However, the amount of penetratin leaving the endo-lysosomal compartment was significantly higher in cells treated with these high atorvastatin concentrations even though the total cellular uptake was lower (Fig. 5). This finding is evidenced by the almost two-times higher NF/AFDye532 intensity ratio characterizing the fraction of penetratin escaping from endosomes. In conclusion, we have convincingly shown that release of penetratin from the endo-lysosomal compartment is the step that is the most significantly increased by atorvastatin treatment.

**Figure 5.**
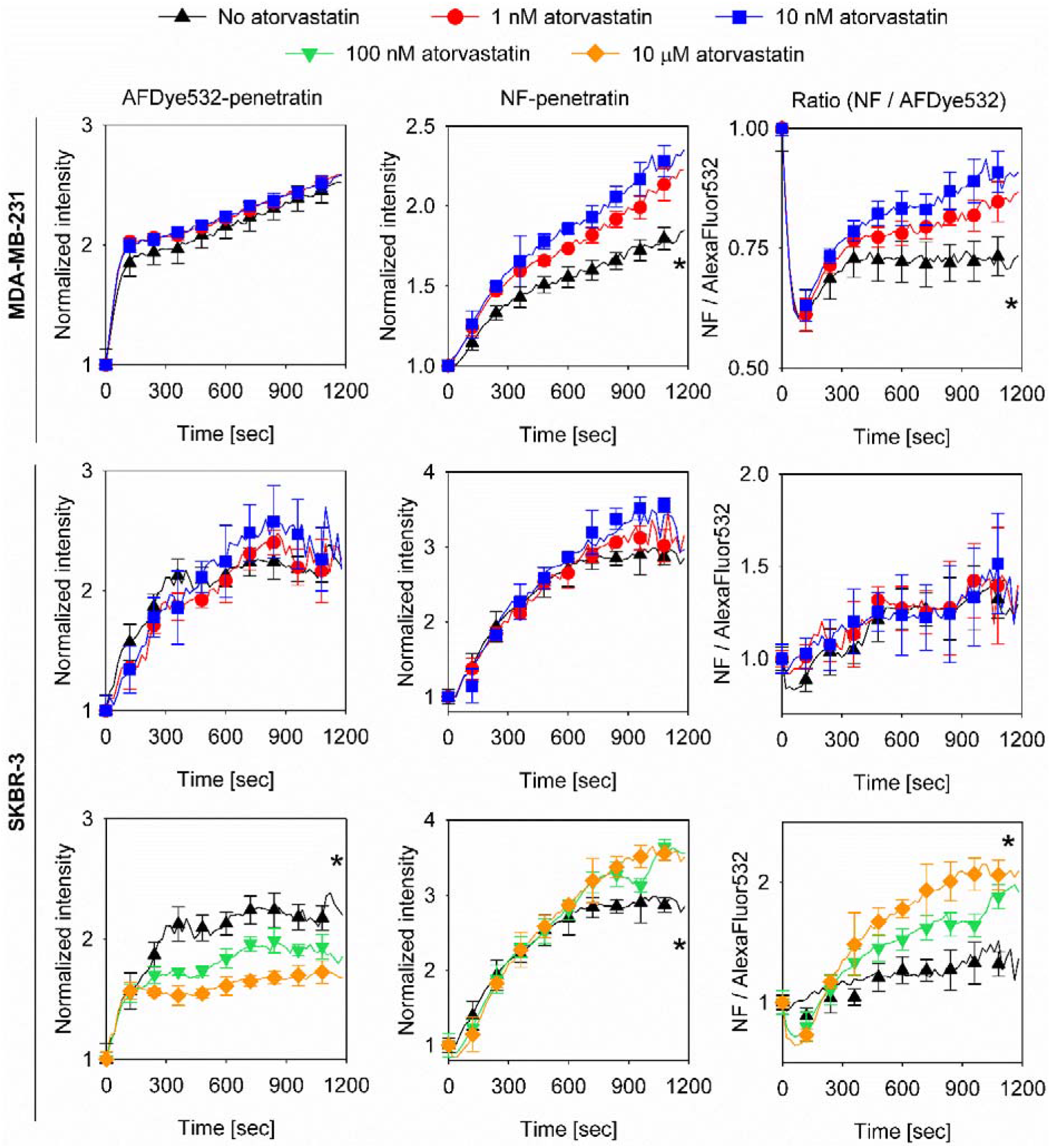
Atorvastatin enhances uptake and endo-lysosomal release of penetratin. Two different cell types were pretreated with the indicated concentrations of atorvastatin for 72 h and the uptake of AFDye-532-penetratin and NF-penetratin were followed by flow cytometry. Time-dependent intensities and their ratio were determined after gating out debris and dead cells. The error bars represent the standard error of the mean calculated from 10-12 samples from four biological replicates. Asterisks indicate significant difference of the sample treated with the highest concentration of atorvastatin in the figure compared to the control at 20 min (p<0.05).

### Characterization of the effects of atorvastatin and the dipole potential modifying compounds on membrane biophysical properties

The previous experiments revealed that treatments reducing the membrane dipole potential enhance the total cellular uptake and the release of penetratin from endosomes, while increasing the dipole potential results in an opposite, weak effect. The validity of this conclusion was checked by calculating the rank-correlation between the dipole potential and penetratin uptake (Suppl. Fig. 4). A significant, negative correlation was revealed between the dipole potential and the total uptake (AFDye532 fluorescence), the concentration of penetratin in non-acidic compartments (NF fluorescence) and the fractional release of penetratin from the endo-lysosomal compartment (ratio of NF and AFDye532 intensities). Since the experimental data corresponding to the effect of phloretin on total cellular uptake of penetratin seemed to deviate from the trend established by the other conditions, the correlation was also tested by removing these outlier data points. The significant and biologically relevant negative correlation between the endo-lysosomal release of penetratin and the dipole potential was retained. However, removing the data points corresponding to phloretin treatment eliminated most of the dipole potential-dependent changes in total cellular uptake (Suppl. Fig. 4). This observation indicates that although both atorvastatin and phloretin decrease the dipole potential, their cellular effects and their influence on penetratin uptake are different. Since our measurement approach based on the pH-dependent fluorescence of NF assumes that the pH of the endo-lysosomal compartment is equally acidic in all compared samples, a possible explanation for the different results obtained with atorvastatin and phloretin is that they alter the pH of lysosomes. Therefore, we performed fluorescence ratiometric measurements of lysosomal pH, which revealed no significant effect of any of the treatments on lysosomal pH (Suppl. Fig. 5). These results imply that the ratio of the fluorescence of AFDye532-penetratin and NF-penetratin correctly reports the release of penetratin from acidic compartments. The issue of the different effect of phloretin and atorvastatin on penetratin release will be further considered in the Discussion.

Since treatments altering the dipole potential could potentially also modify other biophysical properties of the membrane, the correlations between penetratin uptake and two other fluorescent indicators were also tested. Membrane fluidity, inversely proportional to microviscosity and the fluorescence anisotropy of TMA-DPH(50), was not changed by any of the treatments in the SKBR-3 cell line, while it was slightly increased by 6-ketocholestanol in MDA-MB-231 cells (Fig. 3B). In addition, an unexpected increase in the membrane microviscosity of this cell line was induced by atorvastatin, an effect most likely attributable to the compensatory increase in the synthesis of certain lipid species in order to compensate for inhibited cholesterol synthesis (51). Membrane fluidity and total cellular uptake of penetratin did not correlate with each other, but there was a significant correlation between membrane fluidity and the concentration of penetratin in non-acidic compartments (Suppl. Fig. 4). However, membrane fluidity correlated less strongly with the cytoplasmic concentration penetratin than the dipole potential. In addition, the power of membrane fluidity to predict endo-lysosomal escape of penetratin is limited by the fact that none of the treatments induced any significant change in the membrane fluidity of SKBR-3 cells, while phloretin increased endosomal release of penetratin.

The effect of the treatments on the generalized polarization of Laurdan, known to be proportional to membrane compactness (52), was also tested. None of the condition led to any significant change in the membrane compactness in MDA-MB-231 cells, while atorvastatin induces a decrease in this parameter in SKBR-3 cells (Fig. 3C). However, the generalized polarization of Laurdan did not shown appreciable correlation with total penetratin uptake or its concentration in non-acidic compartments (Suppl. Fig. 4). Given the fold-changes induced and the statistical significance of the correlations we can conclude that the dipole potential has the largest power for predicting penetratin uptake, and that release of the cell-penetrating peptide from endosomes and its concentration in the cytoplasm are most strongly correlated with the dipole potential.

## Discussion

In this study correlative measurement of the biophysical properties of the plasma membrane and the uptake of penetratin allowed us to reach the following major conclusions: (i) The physiological, positive membrane dipole potential inhibits the total cellular uptake of penetratin and its release from acidic endo-lysosomes. These conclusions are based on temporal measurement of the cellular intensity of AFDye532-labeled penetratin and the ratio of NF-penetratin to AFDye532-penetratin intensities, respectively. The effect of the dipole potential on both steps of penetratin uptake is most likely mediated by an alteration in the membrane insertion of the cell-penetrating peptide, since incorporation and penetration of peptides and small, hydrophobic molecules into the membrane is known to depend on the dipole potential (27, 53). (ii) Treatment with phloretin and atorvastatin reduced the membrane dipole potential and led to a significantly enhanced cytoplasmic concentration of penetratin. Since the applied, nanomolar concentration of atorvastatin is identical to that used in the clinical setting, the finding of statin-boosted uptake of penetratin is of potential medical significance. (iii) By analyzing the correlation between several membrane biophysical properties and penetratin uptake the dipole potential turned out to be the only characteristic of sufficient value for predicting penetratin uptake.

While both phloretin and atorvastatin decreased the dipole potential and increased penetratin uptake, their differential effects raise question about which step in the uptake process is the most sensitive to the dipole potential. In particular, although atorvastatin decreased the dipole potential more substantially than phloretin, only the latter induces an increase in the total cellular uptake of penetratin, while atorvastatin only increases the fractional escape from acidic endo-lysosomes. These apparent contradictions can be resolved by the following three points: (i) The fact that phloretin does not increase the fraction of penetratin released from the endo-lysosomal compartment is explained by the fact that this compound is unlikely to reach sufficient concentrations in intracellular membranes during the brief, 10-minute incubation and therefore the dipole potential of these compartments remains largely unaffected. On the other hand, the three-day treatment with atorvastatin is sufficiently long so that substantial decrease in the cholesterol content and consequent reduction in the dipole potential of endo-lysosomal membranes can take place. (ii) The minuscule, phloretin-induced decrease of the dipole potential and the significantly elevated uptake of penetratin after phloretin treatment also require clarification. We assume that the physiological level of the dipole potential already limits penetratin uptake as much as potentially achievable by the dipole potential. This conclusion is supported by the fact that the significantly enhanced dipole potential after 6-ketocholestanol treatment had hardly any effect on the characteristics of penetratin uptake. On the other hand, even the relatively minor decrease in the dipole potential, achieved by phloretin, may be sufficient to facilitate penetratin uptake. However, the magnitude of this change in the dipole potential is not sufficiently large to reach statistical significance given the measurement errors. (iii) Although the reduction in the dipole potential achieved by atorvastatin is significantly larger than that of phloretin, atorvastatin had no effect on the total cellular concentration of penetratin. The sudden rise in AFDye532-penetratin intensity and the protracted increase in the intensity of NF-penetratin indicate that initial uptake is mediated by endocytosis, a characteristic left unaltered by any of the treatments. Therefore, the lack of statin-induced increase in the total uptake of penetratin implies that endocytosis must be hindered in atorvastatin-treated samples. Indeed, it has been shown that statins inhibit endocytosis by interfering with the prenylation-dependent function of certain G proteins (54, 55). Alternatively, the compensatory increase in membrane viscosity after atorvastatin treatment may also impede endocytosis.

Despite the aforementioned uncertainties, two of the treatments, phloretin and atorvastatin, significantly enhanced the concentration of penetratin in the compartment most relevant from a therapeutical point of view, the cytosol. Analysis of the correlation between penetratin uptake and different biophysical properties of the membrane revealed that the dipole potential is the strongest predictor of penetratin uptake. Although the dipole potential, membrane compactness and viscosity characterize different membrane properties, all of them are related to membrane order. Therefore, it is not surprising that besides the dipole potential, other membrane characteristics also correlate with penetratin uptake. Albeit in a less consistent way, membrane viscosity is correlated with endo-lysosomal of penetratin. However, the predictive value of this correlation is undermined by the following two points: (i) While phloretin exerts no effect on membrane fluidity, it significantly enhances penetratin uptake in both SKBR-3 and MDA-MB-231 cells. (ii) Although penetratin uptake was modulated by the treatments in SKBR-3, none of them altered membrane viscosity in this cell line. Membrane compactness or hydration, characterized by the generalized polarization of Laurdan, was correlated weakly and in a statistically non-significant way with penetratin uptake. The predictive value of this correlation is further deteriorated by the fact that the generalized polarization of Laurdan was non-significantly modified by any of the treatments in MDA-MB-231 cells, while large and significant effects in penetratin uptake were observed.

In the present manuscript not only did we identify the positive, intramembrane dipole potential inhibiting the uptake and endo-lysosomal release of penetratin, but we could also enhance the cytoplasmic concentration of the cell-penetrating peptide in a medically relevant way by statin treatment. In order for a treatment modality to be of potential medical importance, it must be well-tolerated and therapeutically effective. In contrast to many in vitro studies, in which statins are often overdosed, the nanomolar concentration range of atorvastatin used in the presented experiments is identical to the serum concentration at therapeutic doses (34). By applying atorvastatin not requiring metabolic activation, we also circumvented another pitfall of in vitro cellular studies, the application of statins in a prodrug form, which are unlikely to be converted to the active metabolite in cell cultures. Although certain side-effects are associated with statin treatment, extensive experience indicates that they are safe, well-tolerated even in long-term applications in combinations with other drugs (39, 41). These circumstances further support the potential medical relevance of our findings.

Although reduction of the dipole potential in both MDA-MD-231 and SKBR-3 cells resulted in enhanced penetratin accumulation in the cytosol after phloretin treatment, the latter cell line exhibited lower sensitivity to atorvastatin. Sensitivity to statins correlates inversely with HMG-CoA reductase activity (56, 57). While the expression of this enzyme is higher by ~20-30% in SKBR-3 cells according to a publication (57) and the Expression Atlas of the European Bioinformatics Institute (https://www.ebi.ac.uk/gxa/home), the confidence intervals of the expression of HMG-CoA reductase in the two cell lines completely overlap according to the Genevestigator platform comparing transcriptomic data from several public repositories, with the level of expression corresponding to the MDA-MB-231 cell line spanning three orders of magnitude. In addition, atorvastatin, albeit at a micromolar concentration, resulted in 4-fold induction of HMG-CoA reductase expression in SKBR-3 cells, while no change in enzyme expression was observed in MDA-MD-231 cells (57). Cellular uptake of statins is believed to be mediated by transporters to a large extent. Organic anion transporter polypeptides (OATP) have been implicated in transmembrane import of statins (58–60). However, none of the OATP transporters is expressed significantly differently according to the previous databases. Therefore, the most likely cause for the different atorvastatin sensitivity of the two cell lines in terms of penetratin uptake and reduction in cellular cholesterol content seems to be the difference in the baseline and statin-induced expression of HMG-CoA reductase expression, but solid conclusions cannot be drawn due to inconsistencies in the literature.

In conclusion, we have shown that a decreased, positive membrane dipole potential significantly increases both the total cellular uptake and endocytic escape of penetratin depending on what kind of treatment is used for modifying the dipole potential. As a result, both medically relevant (atorvastatin) and irrelevant (phloretin) treatments decreasing the dipole potential enhance the concentration of penetratin in the cytoplasm, the compartment most relevant for its therapeutic action. This discovery could permit the delivery of drugs and drug candidates exhibiting low or no transmembrane permeability into cells in animal experiments, human trials or in the clinical setting after further studies clarify the cell type dependence and the in vivo potential of this treatment.

## Materials and methods

### Synthesis of penetratin

Penetratin (RQIKIWFQNRRMKWKK-amide, molecular weight 2245.75 g/mol) was synthesized manually on TentaGel R RAM (Rapp Polymere, Tübingen, Germany), a low crosslinked polystyrene PEG copolymer resin (the substitution was 0.2 mmol/g) by the solid-phase method of Merrifield with standard Fmoc chemistry. Amino acids for peptide synthesis were purchased from Iris Biotech (Marktredwitz, Germany). Amino acid side chains were protected as follows: Trt for Gln and Asn, *t*-butoxycarbonyl (Boc) for Trp and Lys, and Pbf for Arg. A typical coupling reaction contained 3 equivalents of Fmoc-amino acid, 3 equivalents of 2-(7-aza-1H-benzotriazole-1-yl)-1,1,3,3-tetramethyluronium hexafluorophosphate (HATU) and 6 equivalents of diisopropylethylamine (DIPEA), and was allowed to proceed with mixing in dimethylformamide (DMF) for at least 2.5 hours. Completion of the coupling was assessed by Kaiser test. The Fmoc group was removed by treatment with 2% piperidine, 2% 1,8-Diazabicyclo[5.4.0]undec-7-ene (DBU) in DMF, followed by six washes with DMF-MeOH-CH2Cl2. Completion of deprotection was assessed by Kaiser test.

### Fluorescence labeling of penetratin

After coupling the last amino acid, arginine, its Fmoc protecting group was replaced by Fmoc-6-aminohexanoic acid in order to introduce a 6-carbon linker followed by removing the Fmoc group. Half of penetratin, still on the resin, was treated with 1.5 equivalents of AFDye532 N-hydroxysuccinimide ester (molecular weight 723.77 g/mol, Fluoroprobes, Scottsdale, AZ) and 1.5 equivalents of DIPEA in DMF overnight. The other half of penetratin was reacted with 1.5 equivalents of 5(6)-carboxynaphthofluorescein *N*-succinimidyl ester (molecular weight 573.51 g/mol, Darmstadt, Germany) in the presence of DIPEA in DMF. Completion of the coupling was assessed by Kaiser test. The labeled peptides were deprotected and released from the resin by treatment with 95:2.5:2.5 (v/v) trifluoroacetic acid (TFA)/triisopropylsilane/water and dithiothreitol for 3 hours. The solutions were filtrated, and the peptides were precipitated with a 15-fold volume of cold diethyl ether. After filtration the crude products were purified by preparative reversed-phase HPLC (JASCO, Victoria, Canada) on a C18 column and lyophilized.

The purity of the products (>95%) was assessed by reversed-phase HPLC (JASCO) equipped with an analytical C18 column. The authenticity of each peptide was confirmed by Bruker electrospray ionization mass spectrometry. The predicted and measured molecular mass of the (M+H)^+^ form of AFDye532-penetratin was 2983.59 and 2983.548, respectively, whereas the predicted and measured molecular mass of the (M+H)^+^ variant of NF-penetratin was 2833.33 and 2832.5, respectively.

### Cell culture

The SKBR-3 cell line was obtained from the American Type Culture Collection (Manassas, VA), while MDA-MB-231 cells were a kind gift of Peter Bay (University of Debrecen). The cell lines were selected based on their different atorvastatin sensitivity (56, 57, 61). Both cell lines were propagated in DMEM containing 10% fetal bovine serum supplemented with antibiotics. Cells were harvested at a confluency of 80-90% before measurements.

### Treatments for modifying the dipole potential or inhibiting HMG-CoA reductase

6-ketocholestanol and phloretin (Sigma-Aldrich, St. Louis, MO) were used for measuring penetratin uptake at increased and decreased dipole potentials, respectively. Both compounds were applied at a concentration of 100 μM at room temperature in the presence of 0.05% (v/v) Pluronic F-127 for 10 min (33, 62). In order to inhibit HMG-CoA reductase atorvastatin (Sigma-Aldrich) was applied at final concentrations of 1 nM-10 μM for 72 hours. The low nanomolar concentrations represent ranges found in human serum during atorvastatin treatment of patients with hypercholesterolemia (34).

### Membrane dipole potential measurement with di-8-ANEPPS

After treatment with atorvastatin, 6-ketocholestanol or phloretin cells were incubated with di-8-ANEPPS (Thermo Fisher, D3167) at a final concentration of 2 μM on ice for 20 minutes (33, 48, 62). Cells were kept on ice until measurement. Right before spectrofluorimetry, the sample was warmed to 37°C, and spectra were acquired with a Fluorolog-3 spectrofluorimeter (Horiba Jobin Yvon, Edison, NJ). The temperature of the cuvette holder was adjusted to 37°C by a circulating water bath. Excitation spectra were registered between 410 nm and 520, while the emission wavelength was set to 660 nm in order to minimize effects of membrane fluidity on the fluorescence of the indicator (62). The ratio of fluorescence intensities integrated between excitation wavelengths 410-416 nm and 503-510 nm correlates positively with the dipole potential (33, 62).

### Measurement of membrane fluidity, hydration and cellular cholesterol content

After treatment with atorvastatin, phloretin or 6-ketocholestanol, 150,000 cells were stained with 10 μM 4’-(trimethylammonio)-diphenylhexatriene (TMA-DPH) or 2 μM Laurdan (6-dodecanoyl-N,N-dimethyl-2-naphthylamine; both obtained from Sigma-Aldrich) for 30 minutes at room temperature. Anisotropy of TMA-DPH was measured with a Fluorolog-3 spectrofluorimeter with the temperature of the cuvette holder adjusted to 37°C. TMA-DPH was excited at 352 nm and its emission was measured at 430 nm. Fluorescence anisotropy (*r*) was measured in the L-format according to the following formula (50):

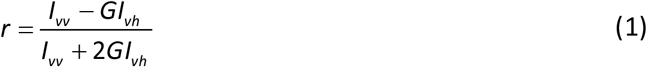

where *I_vv_* and *I_vh_* are the vertical and horizontal components, respectively, of the fluorescence excited by vertically polarized light, and G is an instrument-specific correction factor characterizing the different sensitivity of the detection system for vertically and horizontally polarized light. Laurdan was excited at 350 nm and its emission was detected in the blue range of its emission spectrum between 430-440 nm (*I*_blue_) and at the red edge between 495-505 nm (*I*_red_) to measure generalized polarization *(GP)* according to the following formula (52):

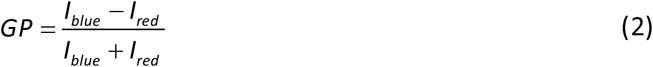

Total cholesterol content of cellular samples was determined using Cholesterol Quantification Kit (MAK043, Sigma-Aldrich).

### Flow cytometric measurement of penetratin uptake and endo-lysosomal release

Cells were incubated in the continuous presence of 5 μM AFDye532-penetratin, 5 μM NF-penetratin and 0.25 μg/ml DAPI at 37°C in the thermostated sample holder of a Becton Dickinson FACSAria III flow cytometer (Becton Dickinson, Mountain View, CA). DAPI was excited at 405 nm, and its emission was measured between 430-470 nm. Both AFDye532 and NF were excited at 561 nm, and their fluorescence was recorded at 515-545 nm and 663-677 nm, respectively. Apart from background measurements every experiment was carried out for 20 min and the measurement started right after adding the DAPI/penetratin mixture, preheated to 37°C, to the cells. After spectral compensation and gating out dead cells based on DAPI positivity the time-correlated fluorescence intensities were exported using FCS Express (De Novo Software, Pasadena, CA). A custom-written Matlab program was used for calculating a moving average with a window size of 20 seconds. The pH-insensitive fluorescence of AFDye532 is proportional to the total cellular uptake of penetratin. Due to quenching of NF at acidic pH, the ratio of NF-penetratin to AFDye532 fluorescence intensities reports the endo-lysosomal escape of the cell-penetrating peptide (47). The fluorescence intensities of both indicators were normalized to the mean intensity measured in the first time window. Normalized mean intensities were divided by each other to obtain the NF-penetratin/AFDye532-penetratin ratio. The standard error of the mean (*SEM*) of the ratio parameter was calculated using error propagation analysis assuming independence of the two fluorescence intensities:

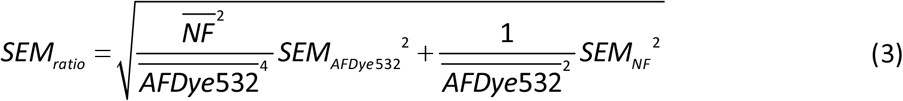

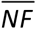 and 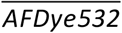 designate the mean intensity of the respective parameter.

### Investigation of penetratin uptake by confocal microscopy

Penetratin uptake and endo-lysosomal release was visualized using a Zeiss LSM880 confocal microscope (Carl Zeiss, Oberkochen, Germany). Fluorescently labeled penetratins were used at the same concentration as in flow cytometry. The temperature of the microscope stage was adjusted to 37°C. AFDye532 and NF were excited at 514 nm and 633 nm, respectively. Detection of AFDye532 and NF fluorescence was carried out between 525-629 nm and 637-759 nm, respectively.

## Acknowledgements

This work was supported by research grants from the National Research, Development and Innovation Office, Hungary (K120302, GINOP-2.3.2-15-2016-00020, GINOP-2.3.2-15-2016-00044). The Lendület grant from the Hungarian Academy of Sciences is gratefully acknowledged. This work was completed in the ELTE Thematic Excellence Program supported by the Hungarian Ministry for Innovation and Technology. Project no. 2018-1.2.1-NKP-2018-00005 has been implemented with the support provided from the National Research, Development and Innovation Fund of Hungary, financed under the 2018-1.2.1-NKP funding scheme.

