## Supplementary material for "Statin-boosted cellular uptake of penetratin due to reduced membrane dipole potential": Suppl.

### Statins boost cellular uptake of penetratin by reducing the membrane dipole potential

**Supplementary figures**

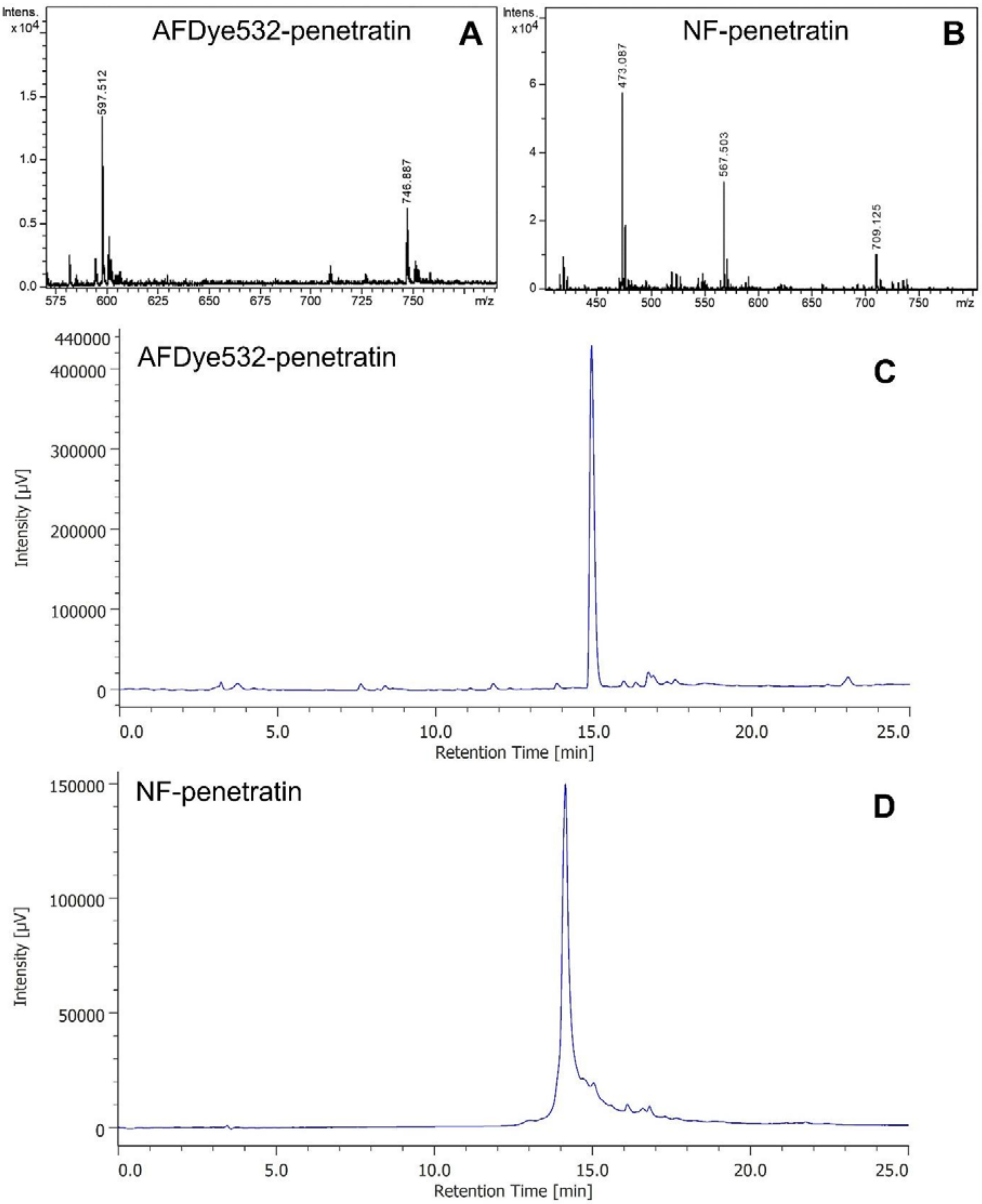


**Supplementary Figure 1.** Mass spectrometric and HPLC analysis of fluorescently labeled penetratin derivatives. Penetratin labeled with AFDye532 or naphthofluorescein (NF) was analyzed by mass spectrometry (A,B) and HPLC (C,D). Electrospray ionization mass spectrometry was performed with a Bruker micrOTOF instrument using a sample concentration of 0.01 mg/ml in methanol. The two peaks in A correspond to the four-times- and five-times-ionized cation of NF-penetratin, whereas the three peaks in B correspond to cations of AFDye532-penetratin with a charge of +4, +5 and +6. Reversed-phase chromatography was performed at a flow rate of 1 ml/min using an injected volume of 20 μl on an analytical column (C18, 250×4.6 mm, particle size: 5 μm, pore size: 100 Å, Phenomenex Luna, 00G-4252-E0). Elution condition: linear AB gradient where A is 0.1% TFA/water and B is 80% acetonitrile/0.1% TFA/20% water. The fraction of B was increased from 5% to 80% in 25 min. Eluted components were quantitated by UV absorption at 220 nm.

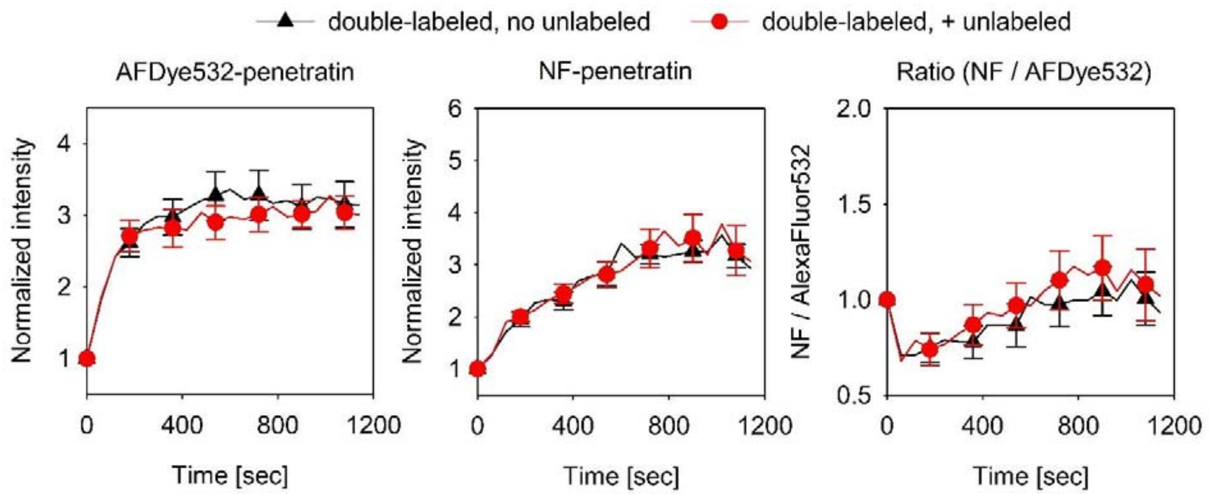


**Supplementary Figure 2.** Resonance energy transfer between AFDye532 and naphthofluorescein does not influence uptake measurements. Cells were incubated with an equimolar mixture of 5 μM AFDye532-penetratin and 5 μM NF-penetratin (black) or with this same mixture of the two fluorescent penetratin derivatives supplemented with 10 μM unlabeled penetratin (red). Cells were continuously kept in the presence of the peptides at 37°C, and the fluorescence intensity of cell-associated fluorescent penetratin was measured by time-correlated flow cytometry. Analysis was performed after gating out debris and dead cells. The error bars indicate the standard error of the mean determined from six samples from two biological replicates.

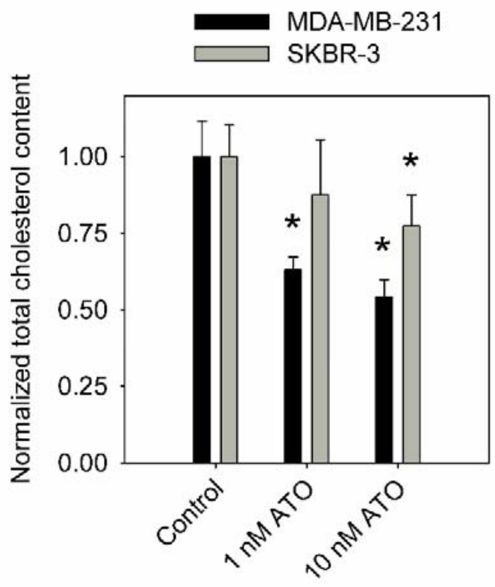


**Supplementary Figure 3.** The effect of atorvastatin on cellular cholesterol content. MDA-MB-231 and SKBR-3 cells were treated with 1 nM or 10 nM atorvastatin (ATO) for three days. Control cells were treated with DMSO. Total (free + esterified) cholesterol content of cells was determined with Cholesterol Quantitation Kit (Sigma-Aldrich, MAK043). The error bars indicate the standard error of the mean determined from seven samples from two independent experiments. Statistical significance was tested by analysis of variance followed by Tukey's HSD test. Asterisks indicate significant ($p<0.05$) difference compared to the control.

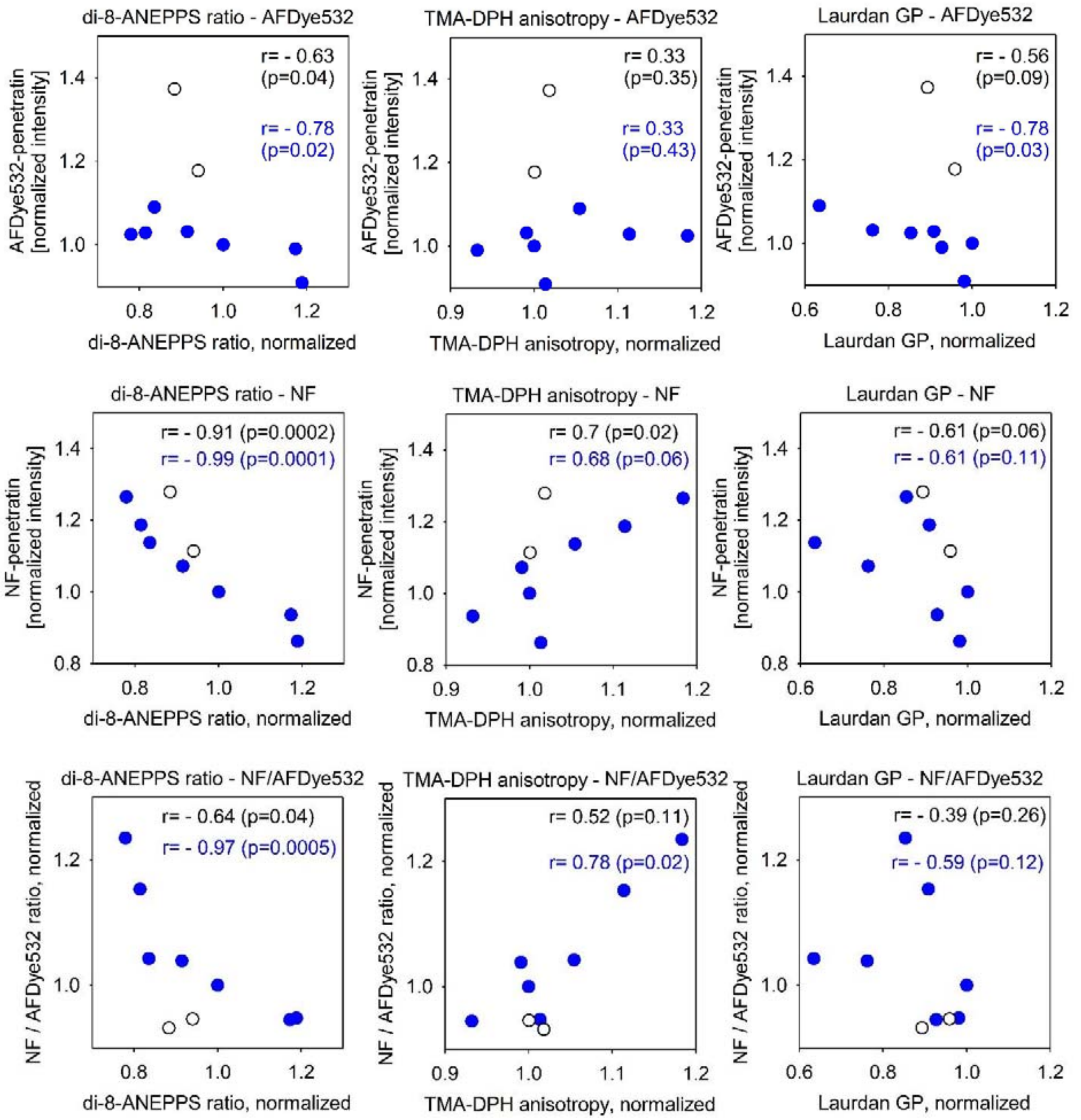


**Supplementary Figure 4.** Correlation between penetratin uptake and biophysical properties of the membrane. The dipole potential was measured using di-8-ANEPPS, whose intensity ratio correlates positively with the dipole potential. Membrane fluidity was assessed using the anisotropy of TMA-DPH correlating inversely with fluidity. The generalized polarization (GP) of Laurdan is inversely correlated with the hydration and positively correlated with the compactness of the membrane. The membrane biophysical properties displayed in these graphs are identical to those shown in Fig. 3, but they were normalized to the control in the case of both cell lines.

The total penetratin uptake, characterized by AFDye532-penetratin fluorescence at 20 min, was double-normalized to the initial intensity at 0 min and to the control sample. Penetratin concentration in the cytoplasm, characterized by NF-penetratin fluorescence at 20 min, was also double-normalized to the initial intensity at 0 min and to the control sample. The fraction release of penetratin from acidic compartments is judged based on the NF/AFDye532 intensity ratio. The white symbols display the results obtained with phloretin in the two cell lines (MDA-MD-231 and SKBR-3), while the blue symbols correspond to data of all other experiments. The Spearman rank correlation coefficient (*r*) and its *p*-value for all data points are shown in black, while the blue text displays the same statistical values for the blue data points only, i.e. for all experiments except those with phloretin.

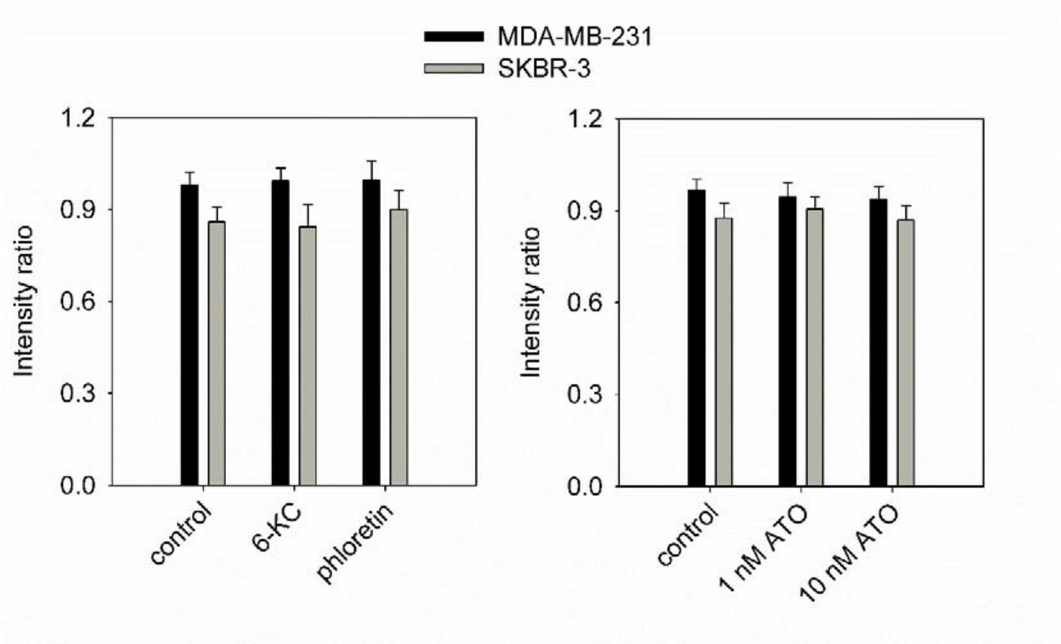


**Supplementary Figure 5.** The lack of any significant effect of the dipole potential-modifying treatments on lysosomal pH. Cells were treated in order to modify the dipole potential as previously, followed by labeling them with 1 μM LysoSensor Yellow/Blue DND-160 (ThermoFisher, L7545) for 10 min. The excitation spectra of the samples was immediately measured by fluorimetry. The emission was measured at 490 nm, and the ratio of intensities excited in the range 305-315 nm and 360-370 nm is shown in the figure. This intensity ratio increases as a function of the lysosomal pH of cells. The error bars represent the standard error of the mean.
